# Disarrangement of Endoplasmic reticulum-mitochondria communication impairs Ca^2+^ homeostasis in FRDA

**DOI:** 10.1101/2020.03.27.011528

**Authors:** Laura R. Rodríguez, Pablo Calap-Quintana, Tamara Lapeña-Luzón, Federico V. Pallardó, Stephan Schneuwly, Juan A. Navarro, Pilar Gonzalez-Cabo

**Author notes:** Equally contribution (Juan A. Navarro, Pilar Gonzalez-Cabo). Corresponding author: Pilar Gonzalez-Cabo.

## Abstract

Friedreich ataxia (FRDA) is a neurodegenerative disorder characterized by neuromuscular and neurological manifestations. It is caused by mutations in gene FXN, which results in loss of the mitochondrial protein frataxin. Endoplasmic Reticulum-mitochondria associated membranes (MAMs) are inter-organelle structures involved in the regulation of essential cellular processes, including lipid metabolism and calcium signaling. In the present study, we have analyzed in both, unicellular and multicellular models of FRDA, an analysis of calcium management and of integrity of MAMs. We observed that function of MAMs is compromised in our cellular model of FRDA, which was improved upon treatment with antioxidants. In agreement, promoting mitochondrial calcium uptake was sufficient to restore several defects caused by frataxin deficiency in Drosophila Melanogaster. Remarkably, our findings describe for the first time frataxin as a member of the protein network of MAMs, where interacts with two of the main proteins implicated in endoplasmic reticulum-mitochondria communication. These results suggest a new role of frataxin, indicate that FRDA goes beyond mitochondrial defects and highlight MAMs as novel therapeutic candidates to improve patient’s conditions.

## Introduction

Friedreich Ataxia (FRDA) is a neurodegenerative disorder mainly caused by homozygous GAA repeat expansion mutations within intron 1 of the *FXN* gene, which encodes frataxin, a protein associated with the mitochondrial inner membrane (Campuzano et al., 1996). The length of GAA expansions decreases the expression of frataxin, which involves neurological and neuromuscular manifestations in patients including progressive trunk and limb ataxia, dysarthria, scoliosis, cardiomyopathy and diabetes mellitus (Parkinson et al., 2013). FRDA is characterized by the degeneration of the large sensory neurons at the dorsal root ganglia (DRG), in charge of proprioception and sense of positioning. This neurodegenerative process affects both, the central and peripheral nervous systems, including spinocerebellar tracts, corticospinal tracts, posterior columns and cerebellum (Koeppen & Mazurkiewicz, 2013).

Frataxin has been proposed to play a role in many physiological functions, mainly related to iron metabolism (Llorens et al., 2019), such as the assembly of the iron-sulfur clusters in the mitochondria, acting as a chaperone (Gerber et al., 2003; Muhlenhoff et al., 2003; Stehling et al., 2004), as an iron-storage protein regulating mitochondrial iron transport (Napier et al., 2005; Yoon et al., 2007) and is associated with heme groups maturation (Schoenfeld et al., 2005; Yoon & Cowan, 2004). Other processes like mitochondrial energy conversion (Ristow et al., 2000) and oxidative phosphorylation (González-Cabo et al., 2005) have been described regarding frataxin, as well as its participation in oxidative stress regulation by reducing the production of reactive oxygen species (ROS)(Napoli et al., 2006; Santos et al., 2010; Tamarit et al., 2016). Moreover, frataxin has an important role in proper calcium (Ca^2+^) handling (Bolinches-Amorós et al., 2014; Mollá et al., 2017), whose effect in neurons is the formation of multiple axonal spheroids mainly caused by Ca^2+^ imbalance.

Mitochondria plays a key role in energy metabolism, specifically in ATP production through oxidative phosphorylation. It is considered the main source of reactive oxygen species (ROS) in most cells, which could generate an imbalance in cellular redox state in circumstances of alteration of the electron transport chain (Chiang et al., 2018). As frataxin is ligated to different mitochondrial processes, its deficiency leads to mitochondrial dysfunction associated with redox imbalance and reduced mitochondrial energy production (González-Cabo & Palau, 2013).

Mitochondrial dysfunction also affects its communication with other cellular compartments and the processes they regulate (González-Cabo & Palau, 2013). In relation to this, we and others have observed increased endoplasmic reticulum (ER) stress in different FRDA models (Bolinches-Amorós et al., 2014; Edenharter et al., 2018). The physical interaction between these compartments termed ER-mitochondria associated membranes (MAMs) (Pinton, 2018) requires specific protein networks. The wide variety of cellular processes in which MAMs are involved in, including lipid metabolism (van Vliet et al., 2014), autophagy (Hamasaki et al., 2013), mitochondrial morphology and cell death (Marchi et al., 2018) are usually altered in several neurodegenerative disorders (Prause et al., 2013; Rodríguez-Arribas et al., 2017; Schon & Area-Gomez, 2013), as well as in FRDA (Abeti et al., 2016; Bolinches-Amorós et al., 2014; Navarro et al., 2010; Simon, 2004).

One of the better characterized functions of MAMs is the rapid exchange of Ca^2+^ between both organelles (Giorgi et al., 2015). The main route for Ca^2+^ stored in the ER into mitochondria is throughout the channels VDAC1/porin and mitochondrial calcium uniporter (MCU), located at the outer and inner mitochondrial membrane, respectively (Britti et al., 2018; Kerkhofs et al., 2018). Ca^2+^ homeostasis is crucial to maintain a proper physiology in different metabolic processes and signaling pathways, specifically mitochondrial regulation of cell survival and oxidative phosphorylation (Berridge et al., 2003). Importantly, we have also previously reported that frataxin-silenced cells show an impairment in Ca^2+^ buffering, as a consequence of reduced mitochondrial Ca^2+^ uptake capacity (Bolinches-Amorós et al., 2014). The involvement of MAMs in FRDA pathophysiology has been recently suggested in a Drosophila model of the disease in which downregulation of fly mitofusin (Mfn) was sufficient to counteract some frataxin-deficient phenotypes via reduction of ER stress (Edenharter et al., 2018). However, undoubtful and definitive evidences regarding the intimate link between frataxin and MAMs are still elusive.

In this work, we assess MAMs architecture and integrity in a well stablished FRDA model in which frataxin is depleted in the SH-SY5Y neuroblastoma cell line using through *FXN* gene silencing two independent short hairpins (FXN-138.1 and FXN-138.2). We confirm that the interactions between the ER and mitochondria are reduced in frataxin-depleted cells, which can be reverted by treating cells with the antioxidant Trolox, a compound that mimics vitamin E. Concomitantly, Trolox also improves mitochondrial Ca^2+^ uptake and alleviates oxidative stress effects in lipid membranes. In agreement with all this, promotion of Ca^2+^ transport into the mitochondria was sufficient to restore several defects triggered by frataxin silencing in the fruit fly, Drosophila melanogaster. Finally, our findings describe for the first time that frataxin is present in MAMs where it interacts with proteins implicated in ER-mitochondria communication highlighting a new role for frataxin in the stability and integrity of these structures. Our results indicate that FRDA pathology goes beyond pure mitochondrial defects and could open new therapeutic avenues for the treatment of the disease.

## Results

### Frataxin deficiency alters MAMs architecture

MAMs encompass a wide variety of proteins, from membrane channels to lipid and glucose metabolism enzymes (Patergnani et al., 2011), but it is mainly mediated by the voltage-dependent anionic channel 1 (VDAC1), located in the mitochondrial outer membrane, the inositol-1,4,5-trisphosphate receptor (IP_3_R) in the ER membrane, and the chaperone glucose regulated protein 75 (GRP75), which links the former two.

To better understand if frataxin levels are able to modify MAMs’ stability or expression, we assessed the connections between the ER and mitochondria in frataxin-depleted cells by performing a proximity ligation assay (PLA) (**Figure 1A-B**) using a neuroblastoma cell line as a model study. Neuroblastoma is a developmental tumor originated from neural crest, like DRG neurons. FXN-138.1 and FXN-138.2 lines were generated through the silencing of the *FXN* gene using two different hairpins, obtaining 82 and 78% of frataxin reduction, respectively (Bolinches-Amorós et al., 2014).

**FIGURE 1:**
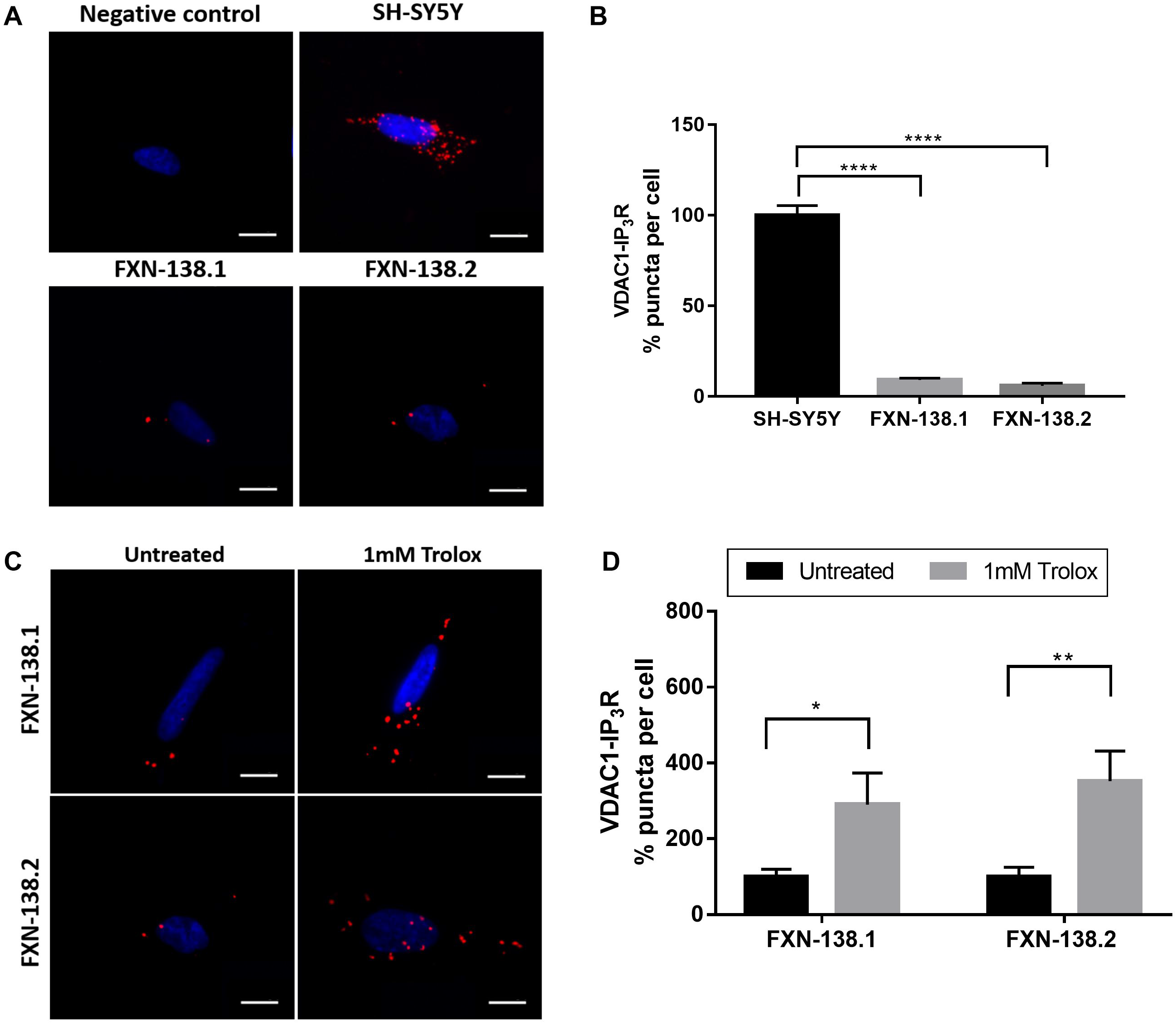
VDAC1-IP_3_R interactions are decreased in frataxin-deficiency cells and can be rescued by Trolox treatment. **(A)** Representative microscopy images of the PLA evaluating the interactions between VDAC1 and IP_3_R in baseline conditions. Red dots represent the physical interactions of the two proteins and nucleus is marked in blue with DAPI. VDAC1 antibody was used as negative control. Scale bars: 12 μm **(B)** Normalized levels of interactions (puncta) per cell in each frataxin-silenced clone compared to SH-SY5Y control. Results (n≥4) are represented as mean ±SEM. A minimum of 500 cells were analyzed in each independent experiment. ANOVA significance: ****p≤0.0001 over SH-SY5Y. **(C)** Representative microscopy images of the PLA evaluating the interactions between VDAC1 and IP_3_R in baseline conditions and after treatments. FXN-138.1 and FXN-138.2 were treated with 1 mM Trolox for 24 hours. **(D)** Normalized number of interactions (puncta) per cell in each frataxin-silenced clone treated compared to untreated conditions. Results (n≥4) are represented as mean ±SEM. A minimum of 500 cells were analyzed in each independent experiment. Student’s t-test significance: *p≤0.05; **p≤0.01 over untreated cells.

Upon frataxin downregulation, we observed a dramatic decrease (around 90%) in the number of interactions between VDAC1 and IP_3_R compared to control cells (SH-SY5Y, 100±5.48%; FXN-138.1, 9.14±1.07%; FXN-138.2, 5.99±1.5%). Accordingly, interactions between VDAC1 and GRP75 were also reduced (SH-SY5Y, 100±2%; FXN-138.1, 26±1.91%; FXN-138.2, 23.78±4.9%) in cells deficient in frataxin (**Figure S1A-B**). Our results indicate that the mitochondrial-endoplasmic reticulum interaction is disturbed in frataxin deficient cells and such impairment could contribute to the mitochondrial Ca^2+^ dysregulation described in frataxin deficiency models, including our previous results (Bolinches-Amorós et al., 2014).

### Trolox recovers MAMs interactions and Ca^2+^ transfer into mitochondria

In order to elucidate the likely relationship between MAMs and Ca^2+^ dyshomeostasis in FRDA, we decided to treat our FRDA cells with the purpose of improving Ca^2+^ management. Trolox is a water-soluble analogue of Vitamin E that acts as lipid peroxidation scavenger and therefore contributes to the stabilization of cellular membranes (Çelik et al., 2018). Interestingly, Abeti and coworkers have recently shown that vitamin E recovers mitochondrial Ca^2+^ uptake in frataxin-depleted cerebellar granule neurons and cardiomyocytes (Abeti et al., 2018b). Thus, we have analyzed the impact of Trolox in the integrity of the ER-mitochondrial contacts. We incubated FXN-138.1 and FXN-138.2 clones with 1 mM Trolox for 24 hours. Importantly, we observed a marked increase (FXN-138.1: untreated 100±19.8%, treated 290.1±83.79%; FXN-138.2: untreated 100±25%, treated 352,2±79.4%) in the number of interactions between VDAC1-IP_3_R (**Figure 1C-D**), which was also confirmed with the VDAC1-GRP75 interactions analysis (**Figure S1C-D**). Our results indicate that Trolox restores communication between these two compartments, probably by reducing oxidative stress environment in mitochondrial and ER membranes.

These results prompted us to analyze in our model the mitochondrial capability of Ca^2+^ uptake after Trolox treatment. FXN-138.2 and control cells were transfected with a mitochondrially targeted chameleon (4mtD3cv) and challenged with 900 μM carbachol in order to release ER Ca^2+^ through IP_3_R (**Figure 2A**). Our results clearly show that SH-SH5Y cells do not show significant changes after Trolox treatment in the area under curve analysis (SH-SY5Y: Untreated 77.46, treated 87.72±3.44). Importantly, the analysis of the mitochondrial response to carbachol showed that trolox successfully boosted Ca^2+^ uptake by mitochondria in cells with frataxin deficiency (**Figure 2B**, FXN-138.2: Untreated 24.59±1.52, Treated 43.78±5.31). These results support the hypothesis that Trolox recovers the network connecting mitochondria and ER and this is sufficient to improve Ca^2+^ transfer between both compartments in frataxin-silenced cells.

**FIGURE 2:**
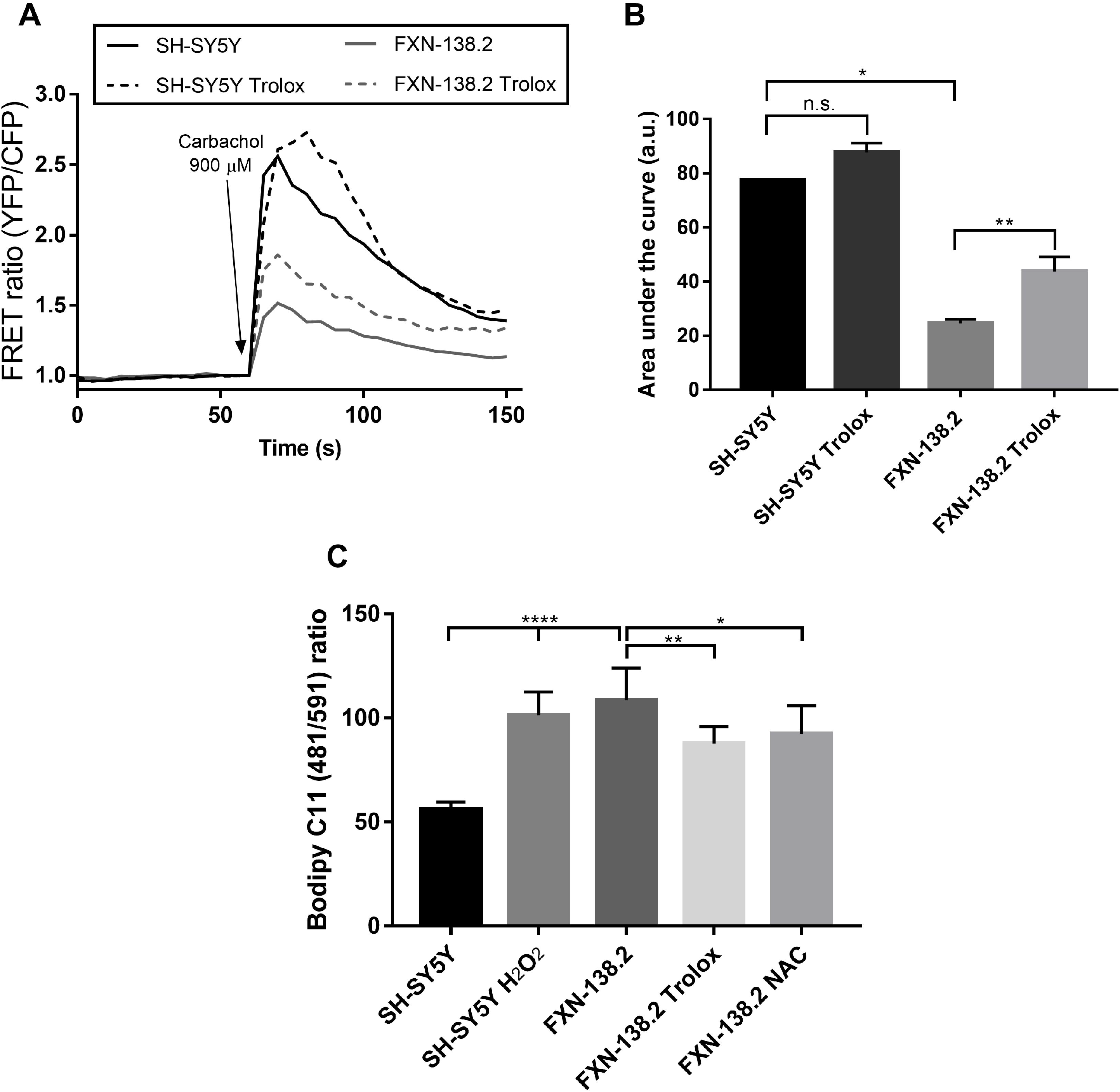
Antioxidants recover mitochondrial calcium uptake and alleviate lipid peroxidation in FXN-138.2 cells. **(A)** FRET analysis of mitochondrial calcium levels measured through 4mtD3cpv chameleon. 900 μM Carbachol was used to stimulate Ca^2+^ release from the ER through IP_3_R channels. Cells were then treated with 1mM Trolox for 24 hours. Results (n≥4) are represented as normalized levels of fluorescence intensity over carbachol loading value. **(B)** Area Under curve (AUC) of the mitochondrial response to carbachol. ANOVA significance: *p≤0.05; **p≤0.01. **(C)** Flow cytometry detection of lipid peroxidation. Cells were stained with 5 μM Bodipy C11 (581/591) for 30 min. Prior to the Bodipy measurements, SH-SY5Y cells were treated with 500 μM H_2_O_2_ as positive control, and FXN-138.2 were treated either with 1mM Trolox or 5mM NAC for 24 hours. Results (n≥4) are represented as 481/591 ratio of the fluorescence intensity of the dye (mean ±SEM). ANOVA significance: *p≤0.05; **p≤0.01; ****p≤0.0001

After confirming Trolox improves ER-mitochondria communication, in both structural and functional ways, we analyzed whether its effects in reducing lipid peroxidation might be underpinning the observed rescue. Quantification of lipid peroxidation by means of Bodipy C11 (481/491) fluorescence intensity by flow cytometry (**Figure 2C**) confirmed a significant increase of these oxidative species in our frataxin-deficient cells. Moreover, we also detected that Trolox was able to alleviate lipid peroxidation in FXN-138.2 cells, suggesting that oxidative insults affect MAMs integrity in FRDA cells.

Furthermore, we also treated FXN-138.2 cells with 5 mM N-acetylcysteine (NAC). NAC is a versatile thiol molecule related to three main antioxidant mechanisms in the cell. It has both direct and indirect antioxidant effects over oxidative species, including the restorage of thiol groups and cysteine donor in Glutathione synthesis, regulating the redox state (Aldini et al., 2018). In agreement with the Trolox results, we observed a recovery of lipid peroxidation levels in frataxin-deficient cells, as well as an increment in the number of VDAC1-GRP75 interactions (FXN-138.2: untreated 100±26%, treated 506.95±100.46%) (**Figure S2A-B**). These results reinforce the idea of an antioxidant-dependent recovery of MAMs and suggest oxidative stress has an important role in Ca^2+^ homeostasis impairment in our FRDA model.

### Frataxin is a member of the protein network of MAMs

Frataxin has been extensively considered as a mitochondrial protein (Schmucker et al., 2008), but interestingly some authors have also described an additional extramitochondrial location (Acquaviva et al., 2005; Condò et al., 2010). Thus, given the reduced levels of MAMs interactions present in our frataxin silenced models, we wondered whether frataxin might be also present at the core of this structures. To elucidate this question, we isolated in our neuroblastoma model the subcellular fractions implicated in the architecture of MAMs (mitochondria, MAMs, ER) and performed a western blot analysis in order to assess the presence of frataxin in the different fractions. Interestingly, we observed that frataxin is clearly located in MAMs, in addition to the characteristic mitochondrial presence (**Figure 3A**). These results indicate that frataxin could have a direct role in the regulation of the contact sites set by mitochondria and the ER.

**FIGURE 3:**
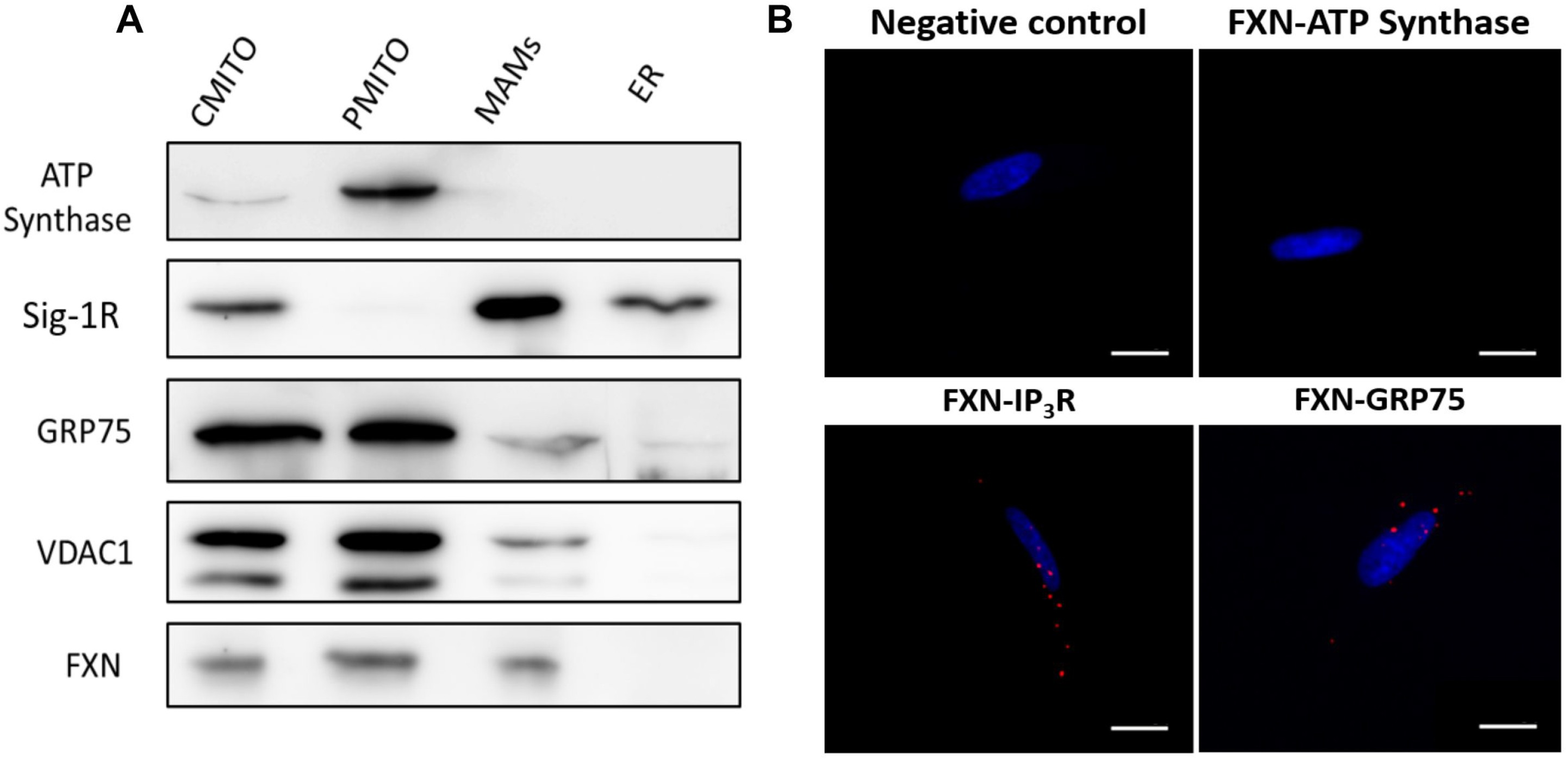
Frataxin is located in MAMs and interacts with two of its main proteins in SH-SY5Y cell line. **(A)** Representative Western blot analysis of frataxin after the isolation of the subcellular fractions implicated in MAMs. ATP Synthase was used as mitochondrial marker. Sig-1R was used as MAMs marker. Other proteins associated to MAMs were used as markers, like GRP75 and VDAC1. Results are representative of at least three independent experiments. CMITO: crude mitochondria fraction (PMITO+MAMs), PMITO: pure enriched mitochondria fraction, MAMs: Mitochondria-ER Associated membranes fraction, ER: endoplasmic reticulum fraction. **(B)** Representative microscopy images of the PLA evaluating the interactions between frataxin-IP_3_R and frataxin-GRP75 in baseline conditions. Red dots represent the physical interactions of the two proteins (either frataxin-IP_3_R or frataxin-GRP75). DNA is marked in blue with DAPI. Frataxin antibody was used as negative control. Interactions frataxin-ATP synthase were evaluated as interaction-negative control. Scale bars: 12 μm

In order to further investigate the possibility of a direct implication of frataxin in the structure of MAMs, we performed a PLA by analyzing the interaction in SH-SY5Y cells of frataxin with GRP75 and IP_3_R, two of the main proteins associated to the mitochondria-ER connections. Importantly, we observed a clear and robust direct relationship of frataxin with both proteins (**Figure 3B**), which suggests a pivotal role of frataxin in the regulation and maintenance of this protein network.

### The induction of ER-mitochondrial contacts recovers brain degeneration in frataxin deficient flies

Next, we decided to evaluate whether the observations obtained in our cellular model might be also relevant in a multicellular organism. We chose a *Drosophila* model in which frataxin silencing was targeted to glia cells because in this model, the interactions ER and mitochondria have been already shown to participate in the development and progression of the pathology (Edenharter et al., 2018). The loss of frataxin in *Drosophila* glia cells using the *Repo-GAL4* line (*Repo*-G4) and a strong RNAi line (*fh*RNAi-1) induces 3 main defects, a locomotor dysfunction, a strong brain vacuolization and the accumulation of lipids within the fly brain (Navarro et al., 2010).

In our first approach, we decided to express the ChiMERA construct in glia cells. The ChiMERA protein is the fusion of the mitochondrial protein Tom70 and the ER protein Ubc6. ChiMERA has been shown to successfully increase mitochondria-ER contacts in yeast and, more importantly, also in flies (Kornmann et al., 2009; Valadas et al., 2018). Then, we assessed whether increasing the bridges between ER and mitochondria was sufficient to recover loss of frataxin in the fly glial cells. Our experiments showed that glial expression of ChiMERA strongly reduced brain degeneration **(Figure 4A-B and Figure 4C)** although no improvement on locomotion **(Figure 4D)** or on the lipid accumulation was observed **(Figure 4A-B)**. Therefore, increasing mitochondrial-ER contacts seems to have a positive albeit minor impact on FRDA phenotypes in glia.

**FIGURE 4:**
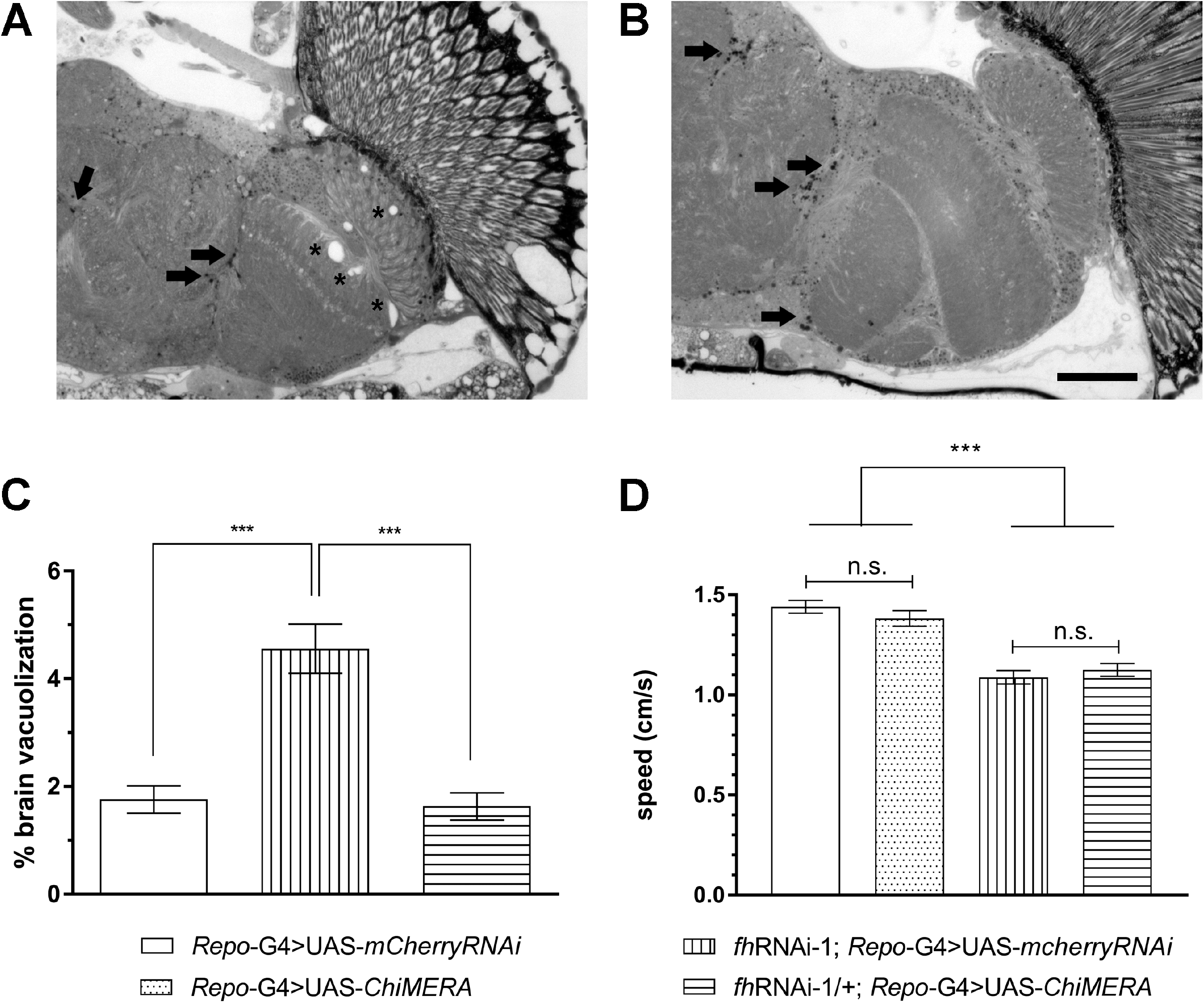
Effects of increasing ER-mitochondrial contacts in frataxin-deficient glia. **(A)** Compared to controls (*Repo*-G4>UAS-*mCherryRNAi*, white column), overexpression of ChiMERA did not alter locomotion in a control background (*Repo*-G4>UAS-*ChiMERA*, dotted column) and failed to restore impaired locomotion of frataxin deficient flies (*fh*RNAi-1;*Repo*-G4>UAS-*ChiMERA*, horizontal stripes). **(B)** Importantly, co-expression of ChiMERA rescued brain degeneration. **(C-D)** Representative microscopy images of semithin plastic sections from brains of frataxin-deficient flies co-expressing either a control UAS (C) or the ChiMERA construct **(D)**. Arrows indicate the presence of the vacuolization that was not observed upon expression of the ChiMERA construct. Graphs represent means ± SEM In A and B data was analysed by One-way ANOVA with *post-hoc* Tukey Multiple Comparison test. ***, P < 0.001, Scale bar: 50 μm.

### Promotion of calcium import into the mitochondria recovers frataxin-deficient phenotypes in Drosophila

As stated in the introduction, frataxin-deficiency has been related to alterations in Ca^2+^ homeostasis in different models, specially a decreased Ca^2+^ buffering by mitochondria, as well as ER stress (Abeti et al., 2018b; Bolinches-Amorós et al., 2014). Furthermore, we have also observed that one of the consequences of reactivating MAMs in our cell culture model was the increase of mitochondrial Ca^2+^ uptake. Therefore, we decided to genetically manipulate Ca^2+^ transport into the mitochondria by altering the expression of the mitochondrial Ca^2+^ uniporter (MCU), since misexpression of MCU or of other components of the MCU complex has been shown to change the mitochondrial Ca^2+^content (Choi et al., 2017; Drago & Davis, 2016; K.-S. Lee et al., 2018; Tufi et al., 2019). We observed that *MCU* mRNA levels were not altered in frataxin deficient flies **(Figure S3A)**. Regarding *MCU* downregulation, unexpectedly a strong silencing using a RNAi line triggered preadult lethality when co-expressed with the *fh*RNAi-1 line, whereas such lethality was not observed in the control cross. To bypass this problem, we decided to use a mutant allele of MCU (MCU1, described in (Tufi et al., 2019)) in a heterozygous configuration. 50% reduction of MCU expression in glia cells did not induce on their own any effect in the three parameters analyzed (negative geotaxis, brain integrity and lipid accumulation, (**Figure S3C-I**), and it is not able to suppress too the locomotor deficit of FRDA flies **(Figure 5A)** and the brain vacuolization **(Figure 5B-E)**. In order to detect the accumulation of lipids *in vivo*, we have used a probe based on a Bodipy fluorophore that allows detection of neutral lipids (see material and methods). As described by Kis and collaborators (Kis et al., 2015) no lipids were detected in control flies **(Figure 5F)**. In agreement with our previous results (Navarro et al., 2010) loss of frataxin in glia cells triggered a strong accumulation of lipids **(Figure 5G)**. However, reduction of MCU expression did not influence the presence of such lipid deposits **(Figure 5H)**.

**FIGURE 5:**
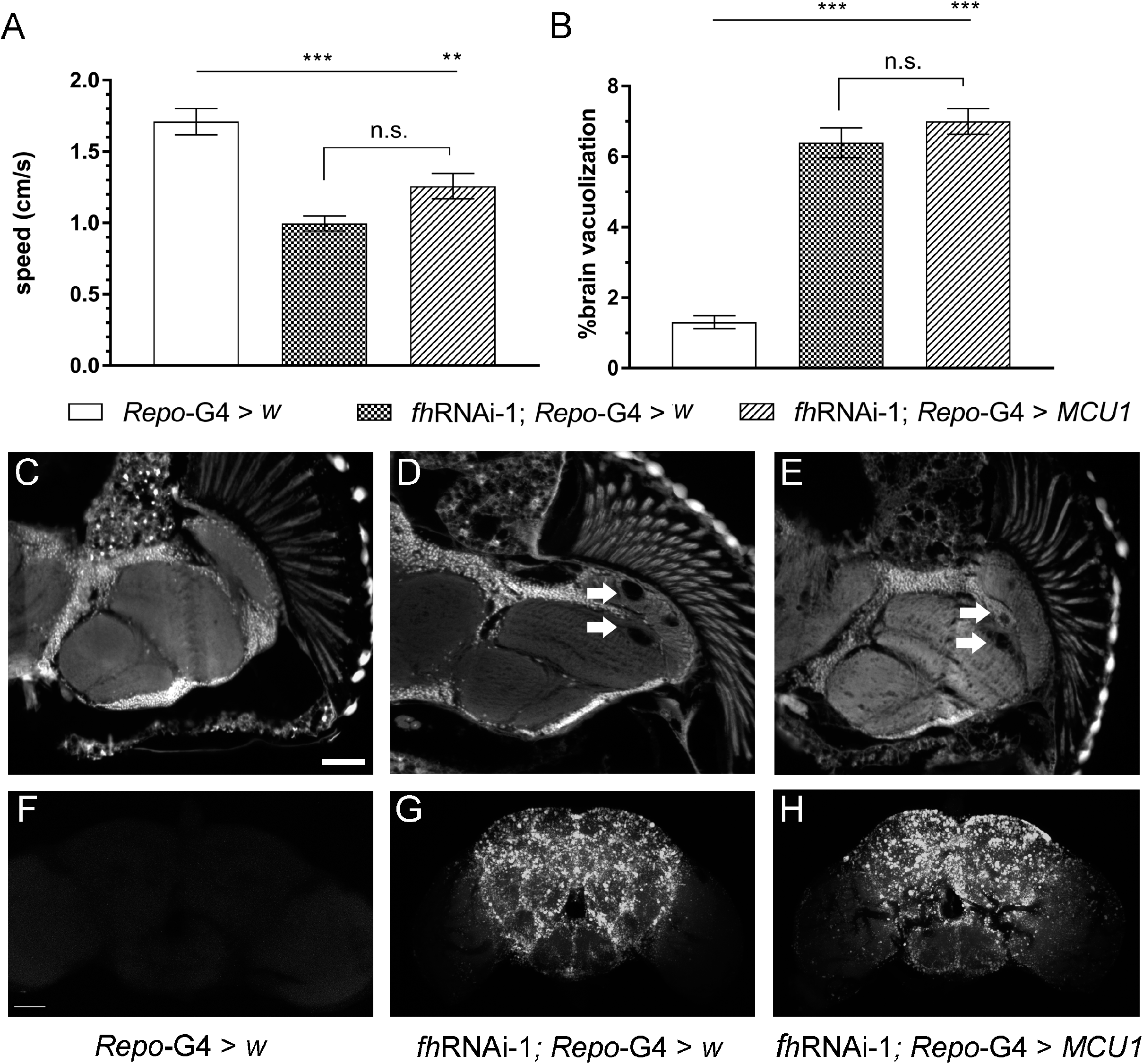
Downregulation of Mitochondrial Calcium Uniporter does not improve loss-of-frataxin conditions. **(A-B)** Compared to control flies (Repo-G4>*w*), downregulation of MCU levels (*fh*RNAi-1;*Repo*-G4>*MCU1*) did not modified the negative geotaxis ability **(A)** as well as the levels of brain degeneration **(B)** of FRDA flies (*fh*RNAi-1;*Repo*-G4>*w*). **(C-E)** Representative paraffin brain sections of controls **(C)**, frataxin-deficient flies **(D)** or the ChiMERA construct **(D)**. Arrows indicate the presence of the vacuolization in frataxin-deficient brains with normal levels of MCU **(D)** and with 50% reduction of MCU expression **(E). (F-H)** Representative Bodipy staining, showing complete absence of lipid deposits in controls **(F)** and a strong accumulation in frataxin deficient flies **(G)** that was not improved by MCU inactivation **(H)**. Graphs represent means ± SEM In A and B data was analysed by One-way ANOVA with *post-hoc* Tukey Multiple Comparison test. **, P < 0.01 ***, P < 0.001, Scale bars: 50 μm.

Next, we decided to increase MCU expression by means of two independent UAS lines (Tufi et al., 2019). Both lines trigger a 10-fold increase of MCU expression **(Figure S3B)**. Interestingly, while promotion of MCU expression in glia cells did not induce any effect in the parameters analyzed in a control background (**Figure S3C-I**), overexpression of MCU completely suppressed the locomotion impairment **(Figure 6A)** as well as the brain vacuolization (**Figure 6B-E**) and the lipid accumulation (**Figure 6F-H**). Since both UAS lines display the same effects in a frataxin-deficient background (as observed in **Figure 6A-B**), only histological findings from line 1 (UAS-*MCUoe1*) are shown and only *UAS-MCUoe1* was used in the resistance to oxidative stress experiments. Reduced longevity under oxidative stress conditions (hyperoxia atmosphere) is another phenotype that is displayed by frataxin deficient flies in glia (Navarro et al., 2010). We confirmed such a defect and observed that, remarkably, overexpression of MCU almost duplicated the median and maximum longevity of FRDA flies under this stressor (**Figure 6I**). Finally, we wanted to assess whether all these rescues of FRDA phenotypes in the *Drosophila* glia were directly linked to an improvement of the mitochondrial function. In this case, we used a second frataxin RNAi line because *fh*RNAi-2 is compatible with a normal development compared to *fh*RNAi-1 when the silencing of frataxin is ubiquitous (*actin-G4*). However, the strong ubiquitous overexpression of MCU achieved with the UAS-MCUoe lines lead to some deleterious effects even in control flies. Therefore, we decided to take advantage of a *Drosophila* Stock containing an EP transposable element in the 5’ untranslated region of the MCU promoter. The EP elements also contain UAS sequences in their structure and therefore they can be used to increase the expression of a nearby gene (Bellen et al., 2004). The EP MCU line only triggers a mild overexpression of MCU (S. Lee et al., 2016). As it can be seen in **Figure 6J**, ubiquitous frataxin knockdown (KD) with *fh*RNAi-2 reduced ATP production around 65% and MCU overexpression was sufficient to restore ATP production in frataxin deficient mitochondria and to trigger a 70% increase of ATP amounts in a control scenario. These results might suggest that lack of mitochondrial Ca^2+^ is also a core component of the FRDA pathology.

**FIGURE 6:**
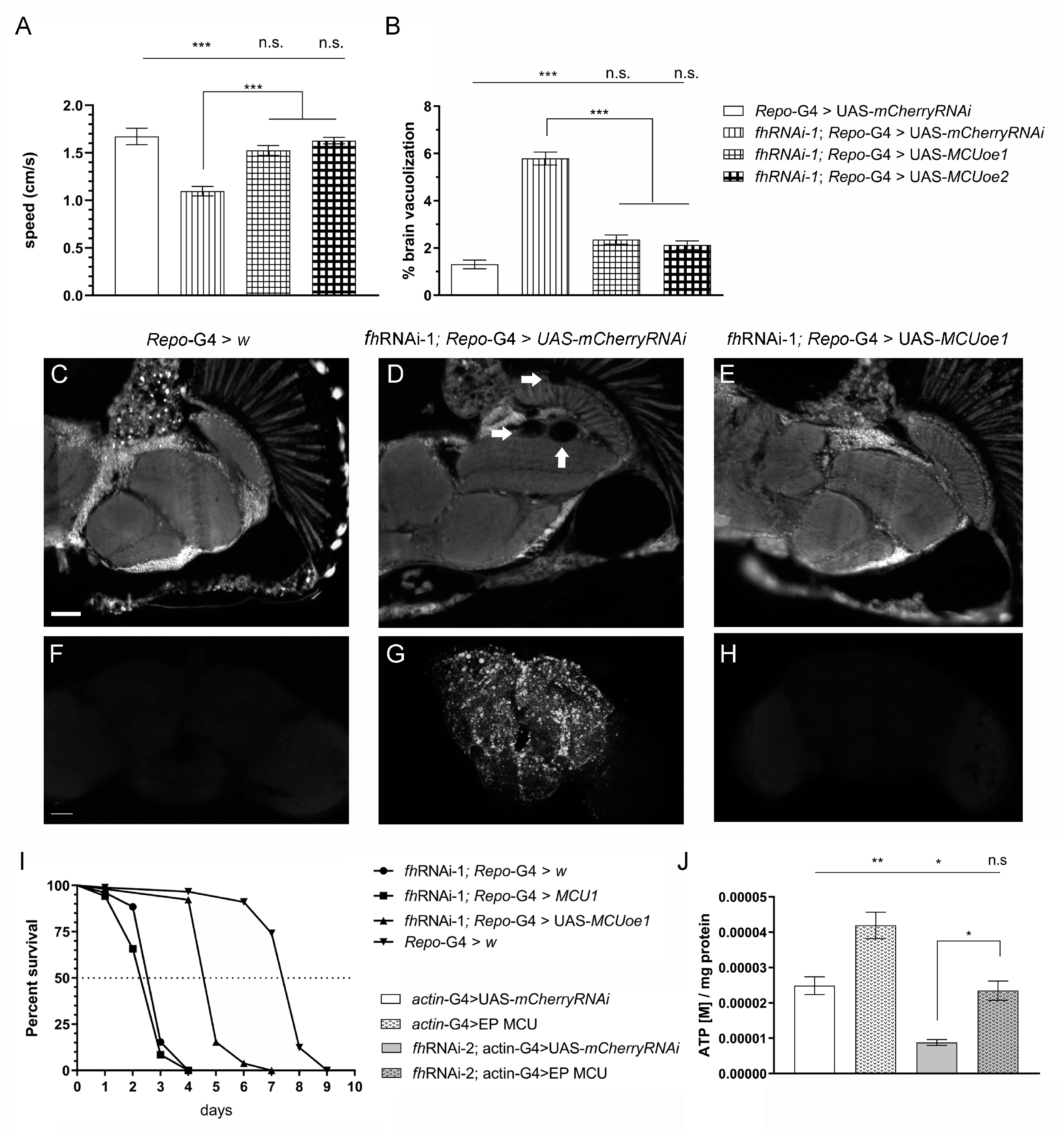
Overexpression of Mitochondrial Calcium Uniporter improves several FRDA-like conditions in flies. **(A-B)** Compared to control flies (Repo-G4>UAS-*mCherryRNAi*, white column), overexpression of MCU along with frataxin silencing flies (*fh*RNAi-1;*Repo*-G4>UAS-*MCUoe1/fh*RNAi-1;*Repo*-G4>UAS-*MCUoe2*) successfully restored to normal values the locomotor performance **(A)** as well as the levels of brain degeneration **(B)** of FRDA flies (*fh*RNAi-*1;Repo*-G4>UAS-*mCherryRNAi*). **(C-E)** Representative paraffin brain sections of controls **(C)**, FRDA flies **(D)** or overexpression of MCU along with frataxin silencing flies **(E)**. Arrows indicate the presence of the vacuolization in frataxin-deficient brains with normal levels of MCU **(D)**. Importantly, increased expression of MCU in glia was sufficient to reverse the degenerative phenotype **(E)** and quantification in **(B)**. **(F-H)** Representative Bodipy stainings, showing complete absence of lipid deposits in controls **(F)**, a strong accumulation in FRDA flies **(G)** and normal lipid levels upon overexpression of MCU along with frataxin silencing flies **(H)**. **(I)** In agreement to our previous results, frataxin silencing in glia reduced longevity under oxidative stress. Importantly, the lack of MCU does not improve survival but MCU overexpression strongly improved this defect. **(J)** Introduction of calcium into the mitochondria, increases ATP values in controls and FRDA flies. Graphs represent means ± SEM In A, B and J data was analysed by Oneway ANOVA with *post-hoc* Tukey Multiple Comparison test. In H, data was analyzed by Log-rank (Mantel-Cox) and Gehan-Breslow-Wilcoxon tests. *, P < 0.05; **, P < 0.01; ***, P < 0.001. Scale bars represent 50 μm

## Discussion

Communication between the mitochondria and the ER is especially important in order to maintain a proper Ca^2+^ transfer required to regulate processes such as energy supply and cell survival. Specifically, there are important physiological functions closely related to mitochondrial Ca^2+^ homeostasis in the nervous system, such as synapse assembly, generation of action potentials and synaptic transmission (Islam, 2017). Variations in the number and structure of MAMs are present in the neurobiology of several neurodegenerative disorders such as Alzheimer’s disease (AD), Parkinson’s disease, Amyotrophic Lateral Sclerosis, hereditary spastic paraplegia and peripheral neuropathies. Interestingly, many proteins involved in these diseases have been found in the ER-mitochondria interface and exhibit an implication the regulation of these structures. For instance, mutations in AD related proteins presenilin 1 and 2 increase ER-mitochondria contacts, upregulating mitochondrial Ca^2+^ uptake. Also, huntingtin, implicated in Huntington’s disease (HD), is essential for ER morphology, so mutated huntingtin activates ER stress and alters Ca^2+^ transfer in MAMs domain (Paillusson et al., 2016; Rodríguez-Arribas et al., 2017). However, little is known about the role of MAMs in the physiopathology of FRDA. In order to address this question, in this work we have performed the first comprehensive analysis of the functionality and integrity of ER-mitochondria contacts also known as MAMs under frataxin-loss conditions.

In the last years we described both in neuroblastoma cells (Bolinches-Amorós et al., 2014) as well as in frataxin deficient sensory neurons of the YG8R mouse model (Mollá et al., 2017) a clear loss of buffering Ca^2+^ capacity in mitochondria leading to increased levels of Ca^2+^ in the cytosol. In agreement, Ca^2+^ chelators were able to improve FRDA conditions (Mollá et al., 2017). More recently, Abeti and co-workers also observed that primary cultures of cerebellar granule neurons and cardiomyocytes from the YG8R mouse model presented a reduced mitochondrial Ca^2+^ uptake that was concomitantly accompanied by lower Ca^2+^ amount in the mitochondria and in the ER (Abeti et al., 2018a). However, the mechanisms underlying an improper mitochondrial Ca^2+^ buffering under frataxin loss conditions have not been studied in detail. It is known that Ca^2+^ is transferred from ER to mitochondria throughout a scaffold formed by IP_3_R, VDAC in the outer mitochondrial membrane (OMM) and the MCU in the inner mitochondrial membrane (IMM). Therefore, alterations in MAMs in frataxin deficient cells would induce a clear Ca^2+^ deregulation. Using a novel experimental approach named proximity ligation assay (PLA) that analyses interactions between proteins, we determined that the number of MAMs was strongly reduced in a neuroblastoma model of FRDA, which correlates with an impairment of Ca^2+^ buffering by the mitochondria.

In order to analyze the impact of MAMs’ dysfunction in the FRDA over Ca^2+^ dyshomeostasis we improved the mitochondrial Ca^2+^ uptake to associate it with a recovery of MAMs interactions. In cardiomyocytes from the YG8R mouse model treated with vitamin E, the mitochondrial capability of Ca^2+^ buffering was restored (Abeti et al., 2018b). Accordingly, treatment with the vitamin E mimic Trolox also improved Ca^2+^ buffering capability in our neuroblastoma model. And more importantly, this was possible by increasing the ER-mitochondria contacts. Thus, the recovery of the network connecting mitochondria and ER is sufficient to improve mitochondrial Ca^2+^ uptake.

In order to test this hypothesis in an *in vivo* model, we decided to take advantage of the outstanding tools for genetic studies offered by the fruit fly (*Drosophila melanogaster*). *Drosophila* FRDA models have provided along the last decade several evidences about FRDA-dependent mechanisms beyond the state of the art (Monnier et al., 2018). Our results with the overexpression of MCU are in favor of the hypothesis that increasing the Ca^2+^ uptake by the mitochondria is neuroprotective in FRDA flies. Using different sensors to visualize and quantify mitochondrial Ca^2+^, overexpression of MCU has been shown to increase basal concentration of mitochondrial Ca^2+^ as well as to enhance Ca^2+^ uptake in drosophila cultured embryonic cells (Chaudhuri et al., 2016) and in living flies (Choi et al., 2017). Therefore, in our experimental scenario, overexpression of MCU would increase the mitochondrial Ca^2+^ content and this would be sufficient to recover several loss-of frataxin phenotypes in the fly. It is known that the cellular bioenergetics are sustained by the transfer of Ca^2+^ from ER to mitochondria via MAMs (Cárdenas et al., 2010). Ca^2+^ seems to influence the activity of some enzymes such as mitochondrial ATP synthase pyruvate dehydrogenase complex (PDC), isocitrate dehydrogenaseand α-ketoglutarate dehydrogenase (Tarasov et al., 2012). All these evidences are validated in our *in vivo* model where we recovered ATP levels due to the reintroduction of Ca^2+^ in the mitochondria.

There are several lines of evidence highlighting the key role of antioxidants in FRDA (Ulatowski & Manor, 2015). In 2008, Cooper *et al*. found promising improvements with a combined Coenzyme Q10-Vitamin E therapy in FRDA patients (Cooper et al., 2008) and NAC, a direct donor of cysteines, has recently been used as a Nuclear factor E2-related factor 2 (Nrf2) modulator in FRDA fibroblasts, exhibiting neuroprotective features and increasing frataxin expression (Petrillo et al., 2019). We have recovered function and structure of MAMs with NAC or Trolox treatment, demonstrating a direct relationship between Ca^2+^ dyshomeostasis and oxidative stress. Several authors have described oxidative stress together with Ca^2+^ dysregulation in different FRDA models (Abeti et al., 2018b; Chiang et al., 2018; Mollá et al., 2019; Wong et al., 1999). It is interesting to emphasize that lipid peroxidation has been described in FRDA patients (Emond et al., 2000) as well as in flies and different tissues from FRDA mice models (Abeti et al., 2015, 2016; Al-Mahdawi et al., 2006; Navarro et al., 2010). Here we show that NAC and trolox treatment decreases membrane lipid peroxidation in our frataxin deficient cell model recovering Ca^2+^ homeostasis. These results underscore the importance of a reduced cytosolic microenvironment surrounding the MAMs to preserve its physiological architecture.

Bringing together these results, we hypothesize frataxin could participate in MAMs in two different but not necessarily exclusive ways. Firstly, our findings describe for the first time frataxin as a member of the protein network of MAMs, indicating a pivotal role in the processes regulated by these structures, including a proper redox environment. Secondly, the fact that frataxin has a direct interaction with GRP75 and IP_3_R, two of the main proteins implicated in mitochondrial Ca^2+^ transfer from the ER, suggesting its implication in the VDAC-GRP75-IP_3_R protein bridge stabilization. These results contribute to understand the mechanisms of Ca^2+^ dysregulation. Our results agree with previous results obtained after artificially increasing the amount of mitochondria-ER connections by means of the expression of the so-called ChiMERA construct (Valadas et al., 2018) in an *in vivo* fly model of FRDA. Genetic manipulation of mitochondrial-ER bridges has been shown to be a successful counteracting strategy in *Drosophila* models of AD (Garrido-Maraver et al., 2020) where the promotion of contacts was beneficial. Here we observed that expression of ChiMERA in glia cells was only able to revert the neurodegeneration whereas locomotion and lipid homeostasis were not improved. Such a limited effect might further support the role of frataxin in the stability of mitochondria-ER contacts, and more concretely in the mitochondrial calcium uptake.

In conclusion, our findings indicate that frataxin deficiency causes an impairment in both the ER-mitochondria communication and in the dysregulation of Ca^2+^ homeostasis, which provides a new approach regarding frataxin functionality and mitochondrial imbalance in FRDA. These results raise the question whether reduction of the interactions between these two organelles is the critical event in the progression of the pathology, offering a new field of investigation regarding MAMs as therapeutic targets.

## Materials and methods

### Cell culture and production of stable SH-SY5Y cell lines

The human SH-SY5Y neuroblastoma cell line was grown in DMEM-12 (Gibco) supplemented with 10% fetal bovine serum and 1% L-glutamine and 1% penicillin/streptomycin. Cells were maintained at 37ºC in an atmosphere of 5% CO_2_ in air. For the generation of stable cell lines with the gene silencing of *FXN*, SH-SY5Y cells were transfected with pLKO.1 vector (MISSION ^®^ shRNA plasmid DNA, Sigma-Aldrich) containing a hairpin sequence of *FXN* (TRCN0000006138) (Bolinches-Amorós et al., 2014). Transfection were performed using SuperFect Transfection (Qiagen) according to the manufacturer’s instructions. The stably transfected cells were selected and maintained in medium with 2 μg/ml puromycin.

### Drosophila Maintenance and Stocks

Fly stocks were maintained at 25ºC on standard cornmeal-agar medium (water: 36 l; yeast: 720 g; cornmeal: 3200 g; soy flour: 400 g; light malt extract: 3200 g; sugar beet syrup: 880 g; agar: 320 g, Nipagin 120 g). The crosses between the GAL4 drivers and the UAS responder lines were carried out at 25ºC. UAS constructs were used in heterozygous configurations for all experiments. Genetic interactions were carried out by mating virgin females from the stocks *fhRNAi-1* / CyO; *Repo-GAL4* / TM6B *tub-GAL80* or *fhRNAi-2* / CyO; *actin-GAL4* / TM6B *tub-GAL80* with males of the corresponding UAS line. The stocks used in this work are described in Table S1.

### Hyperoxia and negative geotaxis assays

In all experiments (normal conditions and food supplementation), flies were raised at 25ºC under a 12hr:12hr light/dark cycle and male individuals were collected within 24-48 hours posteclosion, placed at a density of 25 per vial and transferred to vials with fresh food every 2–3 days. Lifespan experiments were conducted in standard cornmeal agar medium and 100-150 males per genotype were used in each experiment. Hyperoxia treatment started one day posteclosion and was performed by exposing flies in a glass container with a constant flux of 99.5% oxygen under a low positive pressure at 25°C (Botella et al., 2004). Flies were transferred every day to new vials. Locomotor assays were performed as previously described (Botella et al., 2004). 10-12 flies per genotype were assessed and each fly was recorded 3 times and the mean value of each fly was used for the subsequent analysis.

### Semiquantitative Real Time PCR from *Drosophila* Samples

Total RNA was extracted from 15 thoraces using peqGold TriFast reagent (PEQLAB Biotechnologie GMBH, Erlangen, Germany) following manufacturer’s instructions. 500 ng mRNA were converted into cDNA using QuantiTect Rev. Transcription Kit (Qiagen GmbH, Hilden, Germany) and then used for qPCR with ORA qPCR Green ROX L Mix (HighQu, Kralchtal, Germany) on a CFX connect^™^ Real-Time PCR Detection System (Bio-rad, Hercules, California, USA). The ribosomal protein 49 (*rp49*) was used as internal control. The results from at least four independent biological replicates were analyzed with the Bio-Rad CFX manager 3.1 software. Gene expression levels were referred to the internal control, the relative quantification was carried out by means of the ΔΔCt method and the results were plotted as relative mRNA expression. Each experiment consisted of 3-5 independent biological replicates. The genes and primers used for the analysis are: *MCU* - Fw: 5’-CGTCCTGCACCATCGAAAG-3’; Rv: 5’-GTTTGGGAGGATTCACATCCAAT-3’ RP49 - Fw: 5’-CCAAGCACTTCATCCGCCACC-3’; Rv: 5’-GCGGGTGCGCTTGTTCGATCC-3’

### Analysis of Drosophila brain integrity and brain lipid homeostasis

Paraffin sections were performed from 35-day-old adult flies. Flies were fixed with carnoy (ethanol:chloroform:acetic acid at a proportion 6:3:1), dehydrated in ethanol, and embedded in paraffin. Paraffin sections (7 μm) from 10 flies of each genotype were analyzed under a fluorescence microscope. Brain vacuolization was quantified using the Image J 1.48v software. The affected area was referred as % of total brain area that includes the lamina (La), the outer chiasm (OC) and the medulla (Me). For examination of accumulation of lipid droplets, flies of appropriate age and genotype were fixed in 4% PFA for two hours. Brains were then dissected, washed with PBST and incubated 24 h at 4°C with the Bodipy^™^ 493/503 (D3922, Thermo Fischer Scientific, MA, USA) at a dilution 1:500 (from a stock solution 1mg/ml in EtOH). Samples were embedded in Vectashield mounting medium (Vector Laboratories, Burlingame, CA, USA). At least 5-10 flies per genotype were scanned using the Confocal Laser Scanning Platform Leica SP8 (Leica, Germany). Samples were excited with an argon laser at 488nm (20% tube current). Signals were detected using a HC PL APO 40x/1.30 Oil CS2 objective at 490-540nm. Images were generated at a resolution of 1024 x 1024 pixels. Brains were scanned in z-stacks (1 μm) with 30-35 images per brain. All images were acquired using the same exposure, light intensity and filter settings. Confocal images were further processed with the image processing software Fiji 2.0.0 (Schindelin et al., 2012). In detail, background was subtracted in Fiji via the Rolling Ball method (Radius=50 pixels). Maximum projections of 30 slices were made and the resulting image was again subjected to background subtraction. Finally, contrast of each image was adjusted to improve quality of signal. In the experiment expressing the UAS-ChiMERA construct, lipids droplets were studied with light microscopy because the ChiMERA protein is coupled to GFP (Valadas et al., 2018) and this prevents the correct observation of Bodipy^™^ 493/503 signals. In this case, semithin epon plastic sections from 35-day-old adult fly brains were prepared and lipids were identified by characteristically dark droplet-like structures.

### Quantifications of ATP amounts

ATP was determined using ATP Bioluminescence Assay Kit HS II (Roche, Mannheim, Germany) according to manufacturer’s instructions with some modifications. Briefly, five flies were homogenized in 200 μl pre-heated ATP assay buffer (100 mM Tris, 4 mM EDTA, pH 7.75), boiled for two minutes and centrifuged for one minute at 1000g. The supernatant was diluted 1:5 in ATP assay buffer and 50 μl of the dilution were used to measure ATP levels. Luminescence was detected with a Tecan Spark^™^ 10M plate reader (Tecan Trading AG, Switzerland). Each experiment consisted of 3-5 independent biological replicates. ATP quantities were finally referred to protein amount using the Bradford assay (Coomassie Plus^™^ Protein Assay Reagent, Thermo Scientific, Schwerte, Germany).

### Proximity Ligation Assay (PLA)

Cells were seeded in a μ-Dish 35 mm, high Glass Bottom (Ibidi) at a density of 200000 cells/dish. After fixation with 4% paraformaldehyde for 10 minutes, cells were washed with PBS and permeabilized with 0,5% PBS-Triton-X for 10 minutes. Cells were washed with 0,05% PBS-Tween 20 and blocked with Duolink^®^ Blocking solution (Sigma-Aldrich) for 1 h at 37ºC. After incubation, blocking solution was removed and a combination of two of the following specific antibodies were incubated overnight at 4ºC: Frataxin (Thermo Fisher, MA3-085), VDAC1 (Abcam, ab14734), GRP75 (Santa Cruz, sc13967), IP_3_R (Abcam, ab125076), ATP5A (Abcam, ab110273). Cells were incubated at 37ºC for 1 h with Duolink^®^ *in situ* PLA^®^ Anti-rabbit PLUS and Anti-mouse MINUS (Sigma) and later washed with 0,01% PBS-tween. Ligation and amplification steps were performed using Duolink^®^ *in* situ Detection reagents Red (Sigma). Ligase was incubated for 30 minutes at 37ºC and washed with 0,01% PBS-tween. Polymerase was incubated for 100 minutes and washed with 1x and 0,01x SSC buffer respectively for 5 minutes each. Nuclei were detected with DAPI Fluoromount-G^®^ (Southern Biotech). All the washes were performed three times for 5 minutes, at room temperature with agitation. All the incubations at 37ºC were performed in a humidity chamber. Images were obtained using a Leica DMi8 with DC9000GT camera and a 63x oil immersion objective.

### Calcium Imaging

Cells were seeded at a density of 25000 cells/well in a 6-well plate. 16-20 h after, cells were transfected with pcDNA-4mtD3cpv plasmid using Lipofectamine^®^ 3000 Transfection kit (Thermo Fisher). 24 h after transfection, cells were trypsinized and seeded in a μ-Slide 8 Well chamber (Ibidi, 80826), suitable for life imaging. Measurements were carried out 48h after transfection. For all the experiments with carbachol (Merck, 212385), medium was removed prior the beginning of the recordings and replaced with HBSS without Ca^2+^ (Thermo Fisher, 14175095). D3cpv was excited at 435 nm and fluorescence measured at 450-480 nm (CFP) and 520-540 nm (YFP). Images were recorded every 5 seconds using a DC9000GT camera (Leica^®^) and a W-view Gemini (Hamamatsu) for optic splitting imaging. 900 μM Carbachol was added at 60 seconds from the beginning of the experiment. FRET ratio (YFP/CFP) was calculated by measuring fluorescence intensity of the two emission wavelengths, where CFP channel represents fluorophore unbound to Ca^2^ and YFP, Ca^2+^-bond.

### Lipid peroxidation measurements

In order to measure lipid peroxidation, cells were loaded with 5 μM C11 BODIPY (581/591) (Thermo Fisher) for 30 minutes and 4 μg/μL DAPI for 5 min prior to the beginning of the experiments. Fluorescence intensity was detected by flow cytometry with a BD FACSAria^™^ III cytometer equipped with 5 lasers, 355, 405, 488, 561, 640nm (BD, DIVA 6.0 software). Parameters FS-A, FS-H, SS-A were used for cell morphology, lipid peroxidation was measured with C11 BODIPY fluorochrome by selecting the RATIO parameter (586/530 nm) the discrimination of dead cells was performed with DAPI (excitation 405, emission 450/50). 5000 single events (FS-A vs FS-H) were acquired at medium speed. The mean fluorescence of the RATIO parameter of living cells was determined.

### Purification and isolation of mitochondria, MAMs and ER from SH-SY5Y cells

The following protocol is based in the one described in (Annunziata et al., 2015) with some modifications. Cell cultures at 80-90 % confluence were harvested with trypsin and the pellet washed with 1 ml 1x PBS and centrifuged at 700 g for 3 min. The supernatant was removed, and the pellet resuspended in 500 μl of hypotonic buffer (250 mM Sucrose, 20 mM HEPES pH 7.45, 10 mM KCl, 1.5 mM MgCl_2_, 1 mM EDTA, 1 mM EGTA, Protease inhibitors) and kept on ice for 30 min. Cells were then disrupted by passing them through a 30G needle for 20 times. The solution was centrifuged at 700 g for 10 min at 4 ºC and the supernatant transferred to a 2 ml clean tube. The pellet containing nuclei and intact cells was discarded. The collected solution was centrifuged at 10000 g for 20 min at 4 ºC. The supernatant containing the ER was transferred to a clean 2 ml tube and the pellet containing the crude mitochondria was resuspended in 2 ml isolation medium (250 mM mannitol, 5 mM HEPES pH 7.45, 0.5 mM EGTA, 0.1 % BSA). 150 μl of the last suspension was kept as the crude mitochondria (c-mito) fraction. The tube containing the ER was filled with hypotonic buffer, centrifuged at 17000 g for 35 min at 4 ºC, and the new supernatant transferred to an ultracentrifuge tube. The tube was filled with hypotonic buffer and centrifuged at 100000 g for 1 h at 4 ºC. The supernatant was discarded, and the pellet resuspended in 150 μl of hypotonic buffer to obtain the ER fraction. The rest of the solution of isolation medium containing crude mitochondria was transferred to an ultracentrifuge tube filled with 30 % Percoll with gradient buffer (30% Percoll, 225 mM mannitol, 25 mM HEPES pH 7.45, 1 mM EGTA, 0.1 % BSA) and centrifuged at 95000 g for 30 min at 4 ºC. Two bands or fractions are observed at this point, an upper band or light fraction containing the MAMs and a lower band or heavy fraction containing the pure mitochondria. The lower band was collected with a glass Pasteur pipette, diluted in a clean tube with isolation medium and centrifuged at 6300 g for 10 min at 4 ºC. The supernatant was discarded, and the pellet resuspended in 150 μl of isolation medium to obtain the pure mitochondria (p-mito) fraction. The previous upper band containing the MAMs was collected with a glass Pasteur pipette, diluted in isolation medium and centrifuged at 6300 g for 10 min at 4 ºC. The supernatant was transferred to an ultracentrifuge tube filled with isolation medium and centrifuged at 100000 g for 1 h at 4 ºC. The supernatant was discarded, and the pellet was resuspended in 150 μl of isolation medium to obtain the MAM fraction.

### Western Blot Analysis

The isolated ER, c-mito, p-mito and MAM fractions were resuspended in lysis buffer, separated on SDS-polyacrylamide gel, and blotted onto PVDF membranes. Western blots were probed with antibodies against specific protein markers to verify the purity of the fractions and the presence of frataxin. The primary antibodies used were anti-ATP5A (Abcam, ab110273), anti-Sigma1R (Sigma, HPA018002), anti-GRP75 (Sta Cruz Biotec, sc-13967), anti-VDAC1 (Abcam, ab14734) and anti-FXN (Abcam, ab110328). Secondary antibody (anti-mouse or anti-rabbit IgG) coupled to peroxidase was used for detection of the reaction with Amersham ECL Prime. Imaging was performed with an ImageQuant LAS4000 (GE Healthcare).

### Statistical analysis

Survival data were analyzed using the Log-rank (Mantel-Cox) and Gehan-Breslow-Wilcoxon tests. In all further experiments, data is represented as mean ± s.e.m of three to five replicates. When comparing 2 samples, equal variances were confirmed by an F test. When comparing multiples samples, equal variances were confirmed by Bartlett and Brown-Forsythe tests. Normality of data was assessed in all cases with Shapiro & Wilk test and parametric or nonparametric tests were used accordingly. For data that passed normality test, significance was determined by two-tailed T-test or by One-way ANOVA with *post hoc* Dunnett or Tukey Multiple Comparison Test (***, P < 0.001; **, P < 0.01 and *, P < 0.05). Samples that failed normality test were analyzed using non-parametric tests such as Mann-Whitney test or Kluskal-Wallis with *post hoc* Dunn’s Test (***, P < 0.001; **, P < 0.01 and *, P < 0.05). Statistical analysis was carried out using Prism version 8.03 for Windows (GraphPad Software, La Jolla, California, USA, www.graphpad.com).

## Acknowledgements

We would like to thank Gudrun Karch and Ursula Roth for technical assistance as well as Alex Whitworth for the generous gift of MCU fly lines. We also thank the Bloomington Drosophila Stock Center (NIH P40OD018537) and the Transgenic RNAi Project (TRiP) at Harvard Medical School (NIH/NIGMS R01-GM084947) for providing transgenic fly stocks used in this study. This work was supported by grants from the Ministerio de Economía y Competitividad de España [Grant no. SAF2015-66625-R] within the framework of the National R+D+I Plan and co-funded by the Instituto de Salud Carlos III (ISCIII)-Subdirección General de Evaluación y Fomento de la Investigación and FEDER funds; Fundación Ramón Areces (CIVP18A3899); the Generalitat Valenciana (PROMETEO/2018/135); CIBERER (ACCI-2018-22); Joint Research Funding Project from the French National Research Agency (ANR) and the German Research Foundation (DFG) to S.S. (DFG-SCHN 558/9-1). CIBERER is an initiative developed by the Instituto de Salud Carlos III in cooperative and translational research on rare diseases.

## Author contributions

LRR. conducted and designed experiments, analyzed the results and wrote the manuscript. PC-Q and TLL. performed experiments. FVP and SS. interpreting the data and wrote the manuscript. JAN and PG. designed the study, supervised the experiments, analyzed the data and wrote the manuscript. All authors read and approved the final manuscript.

---

**Table S1.** Drosophila melanogaster stocks used in this work

| Name | Genotype | Kindly Provided |
| --- | --- | --- |
| **W1118** | w[1118] | Bloomington Stock 3605 |
| ***actin*-GAL4** | y[1]w[*];P{w[+mC]=Act5C-GAL4}17bFO1/TM6B, Tb[1] | Bloomington Stock 3954 |
| ***Repo*-GAL4** | w[1118]; P{w[+m*]=GAL4}repo/TM3, Sb[1] | Bloomington Stock 7415 |
| ***Mef2*-GAL4** | y[1] w[*]; P{w[+mC]=GAL4-*Mef2*.R}3 | Bloomington Stock 27390 |
| ***Elav*-GAL4, UAS-*Dicer2*** | P{w[+mW.hs]=GawB}elav[C155] w[1118];<br>P{w[+mC]=UAS-*Dcr-2*.D}2 | Bloomington Stock 25750 |
| **UAS-*mCherryRNAi*** | w[1118]; P{w[+mC]=UAS-*fhRNAi-1*} | Bloomington Stock 35787 |
| ***fhRNAi-1*** | w[1118]; P{w[+mC]=UAS-*fhRNAi-1*} | Prof. John P. Phillips |
| ***fhRNAi-2*** | y[1] w[*]; P{w[+mC]= UAS-*fhRNAi-2*}2 | Prof. Maria D. Moltó |
| **UAS-*fh*** | y[1]w[*];P{w[+mC]=UAS- *fh*}2 | Prof. Maria D. Moltó |
| **UAS-*mitoGFP*** | P{w[+mC]=UAS-mito-HA-GFP.AP}/CyO | Bloomington Stock 8442 |
| **UAS-*MCURNAi* (TRiP)** | y<sup>1</sup> sc<sup>*</sup> v<sup>1</sup> sev<sup>21</sup>; P{TRiP.HMS05618}attP40 | Bloomington Stock 67857 |
| ***MCU1*** | w[1118]; MCU1/TM6B, P{w[+mW.hs]=Ubi-GFP.S65T}PAD2, Tb[1] | Prof. Alex Whitworth |
| **UAS-*MCUoe1*** | y[1]w[*];P{w[+mC]=UAS-MCU-flag}attP40 | Prof. Alex Whitworth |
| **UAS-*MCUoe2*** | y[1]w[*];P{w[+mC]=UAS-MCU-flag}attP2 | Prof. Alex Whitworth |
| **EP MCU** | y[1] w[67c23]; P{w[+mC] y[+mDint2]=EPgy2}<br>MCU[EY08610] | Bloomington Stock 19933 |
| **UAS-*ChiMERA*** | y[1] M{vas-int.Dm}ZH-2A w[*];P{w[+mC]=UAS-*ChiMERA*}VK00037 | Prof. Patrik Verstreken |

**Figure S1: VDAC-GRP75 interactions are decreased in frataxin-depleted cells and can be restored by Trolox treatment. (A)** Representative microscopy images of the PLA evaluating the interactions between VDAC1 and GRP75 in baseline conditions. Red dots represent the physical interactions of the two proteins and nucleus is marked in blue with DAPI. VDAC1 antibody was used as negative control. Scale bars: 12 μm **(B)** Normalized levels of interactions (puncta) per cell in each frataxin-silenced clone compared to SH-SY5Y control. Results (n≥4) are represented as mean ±SEM. A minimum of 500 cells were analyzed in each independent experiment. ANOVA significance: ****p≤0.0001 over SH-SY5Y. **(C)** Representative microscopy images of the PLA evaluating the interactions between VDAC1 and GRP75 in baseline conditions and after treatments. FXN-138.1 and FXN-138.2 were treated with 1 mM Trolox for 24 hours. **(D)** Normalized number of interactions (puncta) per cell in each frataxin-silenced clone treated compared to untreated conditions. Results (n≥4) are represented as mean ±SEM. A minimum of 500 cells were analyzed in each independent experiment. Student’s t-test significance: *p≤0.05; over untreated cells.

**Figure S2: N-acetyl cysteine restores the number of connections ER-mitochondria in frataxin-deficiency FXN-138.2 cells. (A)** Representative microscopy images of the PLA evaluating the interactions between VDAC1 and GRP75 in baseline conditions and after treatments. FXN-138.1 and FXN-138.2 were treated with 5 mM N-acetyl cysteine (NAC) for 24 hours. Red dots represent the physical interactions of the two proteins and nucleus is marked in blue with DAPI. Scale bars: 12 μm **(B)** Normalized number of interactions (puncta) per cell in each frataxin-silenced clone treated compared to untreated conditions. Results (n≥4) are represented as mean ±SEM. A minimum of 500 cells were analyzed in each independent experiment. Student’s t-test significance: *p≤0.05; over untreated cells.

**Figure S3: Impact of MCU manipulation in glia cell on fly behavior, brain integrity and lipid metabolism. (A)** MCU levels are not modified upon frataxin downregulation ubiquitously in the whole fly or specifically in fly muscles or neurons. **(B)** Compared to control flies, UAS *MCU* lines are able to upregulate it by ten times, whereas EP MCU line only upregulates MCU expression by 2-fold. **(C)** Nor *MCU* downregulation neither targeted overexpression of MCU in glia impair locomotor activity in Drosophila. **(D-E)** Representative paraffin brain sections of controls **(D)**, MCU-deficient flies **(E)** and flies with a targeted glial expression of MCU **(F)**. **(F-H)** Representative Bodipy stainings, showing complete absence of lipid deposits in controls **(G)** as well as in flies with lower **(H)** of higher **(I)** levels of MCU expression in glia cells. Graphs represent means ± SEM. In A, data was analysed by two-tailed T-test, whereas in B and C, data was analysed by One-way ANOVA with *post-hoc* Tukey Multiple Comparison test. ***, P < 0.001. Scale bars represent 50 μm.

